# Tumor-derived mitochondria enhance CD8^+^ T cell cytotoxicity through SPHK2-dependent S1P signaling

**DOI:** 10.64898/2026.05.15.725367

**Authors:** Chuanfang Chen, Xiunan Wang, Haige Li, Qingxiang Gao, Jia Zhang, Shih-Chin Cheng

**Affiliations:** State Key Laboratory of Cellular Stress Biology, School of Life Science, Faculty of Medicine and Life Sciences, Xiamen University, Xiamen 361102, China; Department of Gastroenterology, Zhongshan Hospital, School of Medicine, Xiamen University, Xiamen, Fujian, China; Department of Internal Medicine, Radboud university medical center, 6500, HB, Nijmegen, the Netherlands

**Keywords:** Mitochondrial transfer, CD8^+^ T cell, S1P-S1PR1, Cytotoxicity

## Abstract

Intercellular organelle exchange is increasingly recognized as a feature of the tumor microenvironment, but whether tumor-derived mitochondria functionally shape anti-tumor T cell immunity remains unclear. Here we show that tumor-infiltrating CD8^+^ T cells acquire functional mitochondria from tumor cells through a contact-dependent, TCR-independent mechanism requiring the mitochondrial trafficking machinery Trak1–Miro1. Transferred tumor mitochondria enhanced CD8^+^ T cell effector activity, increasing cytotoxic molecule expression and tumor-cell killing. Mechanistically, tumor-derived mitochondria carried sphingosine-1-phosphate (S1P), which engaged S1PR1 signaling in recipient T cells. Tumor-specific deletion of Sphk2 diminished mitochondrial transfer-associated T cell activation, impaired CD8^+^ T cell effector function, and accelerated tumor progression in vivo. These findings reveal tumor-to-T cell mitochondrial transfer as an unexpected immunostimulatory circuit in the TME and identify mitochondrial SPHK2–S1P signaling as a regulator of CD8^+^ T cell anti-tumor function.

## Introduction

Horizontal mitochondrial transfer (HMT), the intercellular shuttling of mitochondria, has emerged as a fundamental mechanism of metabolic crosstalk, enabling cells to adapt to physiological stress and environmental cues^1–5^. Within the metabolic desert of the tumor microenvironment (TME), HMT is particularly consequential for CD8^+^ T cell fitness, though its functional impact remains a subject of intense debate. Current models primarily describe a “hijacking” relationship where tumor cells actively deplete mitochondria from infiltrating T cells via tunneling nanotubes, thereby fueling tumor bioenergetics and driving T cell exhaustion^6^. Conversely, the acquisition of tumor-derived mitochondria by T cells has been linked to immune suppression when the transferred organelles carry deleterious loads, such as mutant mitochondrial DNA that impairs respiratory capacity^7^.

However, this “suppressive” view of HMT is challenged by evidence from non-malignant contexts, where bone marrow-derived mesenchymal stem cells (BMSCs) can “recharge” CD8^+^ T cells via mitochondrial donation, effectively bolstering their anti-tumor capacity^8^. These divergent outcomes suggest that the functional fate of a recipient T cell is determined not merely by the acquisition of the organelle, but by the specific metabolic signaling cargo sequestered within the transferred mitochondria.

Sphingosine-1-phosphate (S1P) is a key bioactive lipid mediator generated by sphingosine kinases 1 and 2 (SPHK1 and SPHK2)^9^. The subcellular localization of S1P production is enzyme-specific: SPHK1-derived S1P is primarily cytoplasmic, whereas SPHK2 generates S1P in the nucleus, endoplasmic reticulum, and mitochondria^10–12^. Signaling through a family of Sphingosine-1-phosphate receptors (S1PR1–5), S1P critically regulates lymphocyte migration and T cell function^13,14^. Within the tumor microenvironment, the S1P-S1PR1 axis has been reported to exert both pro-tumor effects—such as promoting tumor cell proliferation and the recruitment of Treg cells^15,16^—and anti-tumor effects, such as normalizing tumor vasculature to enhance anti-tumor therapies^17^. While the S1P-S1PR1 axis is known to influence T cell residency and vascular normalization in tumors, whether HMT can serve as a vehicle for the direct intercellular delivery of mitochondrial-tethered S1P remains unknown.

Here, we demonstrate that tumor-infiltrating CD8^+^ T cells selectively acquire intact mitochondria from MC38 and B16-F10 cells. This acquisition is mediated by a TCR-independent, Trak1–Miro1-dependent transport mechanism. In contrast to the exhaustion-associated “hijacking” model, we find that this transfer directly enhances T cell cytotoxicity. We show that tumor-derived mitochondria act as metabolic “Trojan horses” that deliver SPHK2-generated S1P to the recipient T cell. This localized S1P pool triggers S1PR1 signaling and receptor internalization, providing a requisite signal for increased Granzyme B and Perforin production. Our findings redefine HMT as a positive regulator of anti-tumor immunity and identify the mitochondrial S1P-S1PR1 axis as a critical checkpoint in myeloid-T cell metabolic communication.

## Results

### Tumor-infiltrating CD8^+^ T cell acquire mitochondria from tumor cell

Because lipophilic dyes often leak and produce artifacts in monitoring mitochondrial transfer^18^, we generated B16 melanoma cells stably expressing a mitochondria-targeted fluorescent protein (Cox8a-GFP) to investigate intercellular mitochondrial transfer (Fig. S1A). Following subcutaneous inoculation of Cox8a-GFP B16 cells into mice, we used flow cytometry to evaluate mitochondrial acquisition by tumor-infiltrating immune cells (Fig. 1A). Our results revealed that a substantial proportion of tumor-associated macrophages, neutrophils, and dendritic cells acquired the tumor-derived mitochondrial GFP signal (Fig. S1B–D). To exclude the possibility of artifacts driven by the Cox8a peptide, the GFP tag, or the B16 background, we engineered MC38 colon carcinoma cells to express alternative mitochondrial markers (Cox8a-BFP or Tomm20-GFP). These markers strictly co-localized with MitoTracker Red (Fig. S1A). Upon subcutaneous inoculation of these MC38 lines into C57BL/6 mice, tumor-infiltrating myeloid cells similarly acquired the mitochondrial signals, with GFP^+^ macrophages reaching 70–90% of the total population (Fig. S1D-F). Given that macrophages are highly phagocytic^19^, we reasoned that the high percentage of acquired GFP signal might result from phagocytic uptake of tumor cell rather than functional mitochondria transfer. If mitochondria are phagocytosed, they should be delivered to lysosomes for degradation^20^; conversely, if they are functionally transferred, they should integrate into the endogenous mitochondrial network. Indeed, imaging of sorted tumor-infiltrating macrophages co-stained with MitoTracker Red and LysoTracker demonstrated clear co-localization of the tumor-derived GFP⁺ signal with lysosomes (Fig. S1G). This result confirms that macrophages acquire tumor mitochondrial GFP signal primarily through phagocytic uptake.

**Fig. 1.**
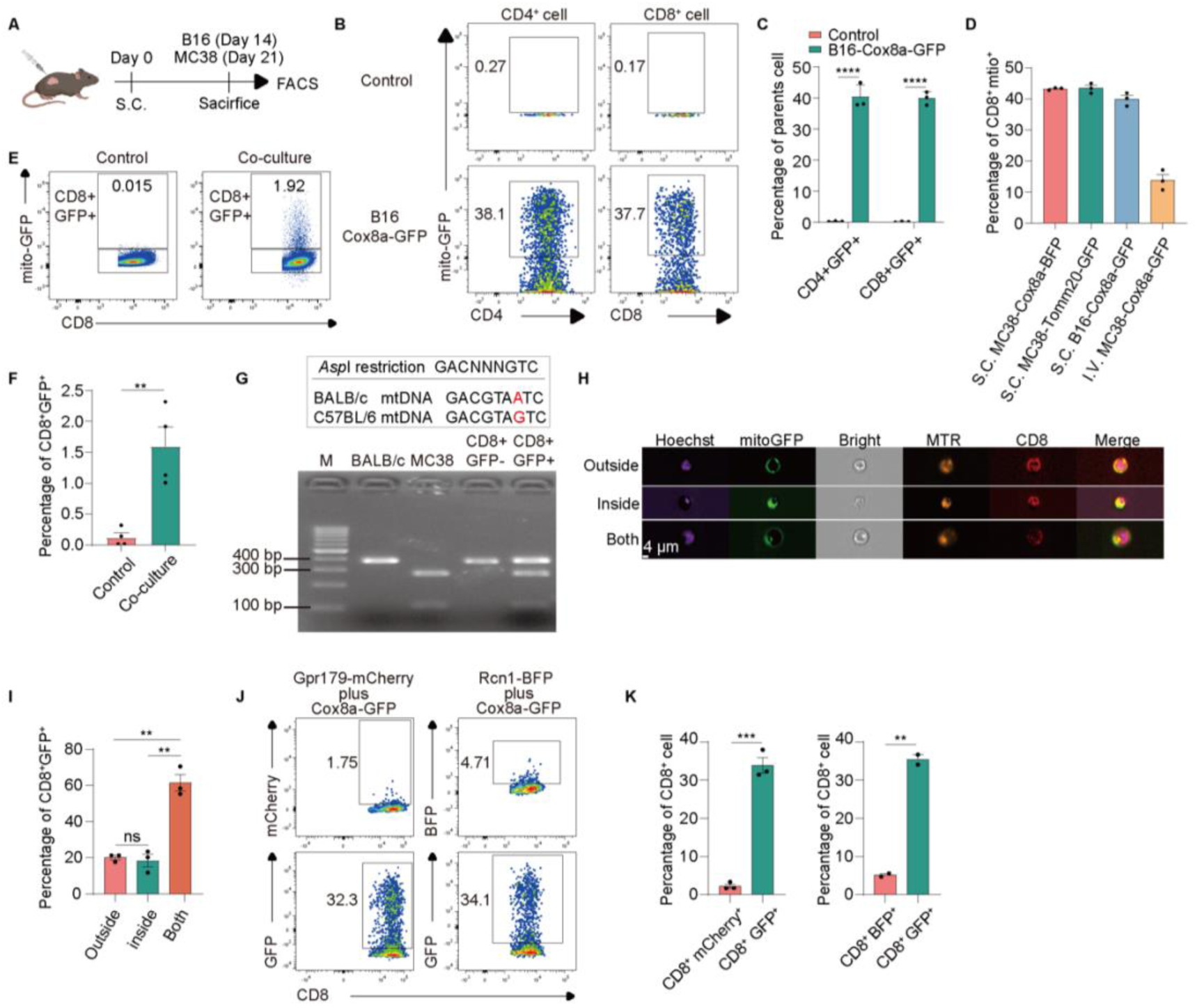
Tumor cell transfer mitochondria to T cell. **A**, Schematic Diagram of the Experiment. **B-C**, Graphical images (**B**) and quantitative analysis (**C**) of T cells acquiring tumor mitochondria *in vivo*. **D**, Quantitative analysis of CD8⁺ cells acquiring tumor mitochondria *in vivo* across tumor type. **E-F**, Representative image (**E**) and quantitative analysis (**F**) of CD8⁺ cells acquiring tumor mitochondria *in vitro* co-culture system. **G**, PCR detection of *mt-Co3* sequence in MC38 tumor-infiltrating CD8⁺ T cells from BALB/c mice. **H-I**, Representative image (**H**) and quantitative analysis (**I**) of CD8⁺ cells acquiring tumor mitochondria *in vivo*, assessed by imaging flow cytometry. **J-K**, Graphical representation and analysis of *in vivo* acquisition of fluorescently-labeled organelles by tumor-infiltrating CD8⁺ T cells, shown schematically in (**J**) and quantified in (**K**). None Significance: ns (*P* > 0.05), \**P* < 0.05, \*\**P* < 0.01, \*\*\**P* < 0.001, \*\*\*\**P* < 0.0001 (unpaired Student’s t-test).

While T cells do not perform classical phagocytosis, both tumor-infiltrating CD4^+^ and CD8^+^ T cells acquired the mitochondria-targeted GFP signal from B16 tumors (Fig. 1B-C). Given their central role in anti-tumor immunity, we focused our investigation on CD8⁺ T cells. We verified the generality of this acquisition by inoculating C57BL/6 mice with MC38 cells expressing alternative markers (Cox8a-BFP or Tomm20-GFP), as well as B16-Cox8a-GFP cells. In all models, including both subcutaneous and intravenous inoculation, tumor-infiltrating CD8^+^ T cells acquired the tumor-derived mitochondrial signal (Fig. 1D). Furthermore, CD8⁺ T cells co-cultured with MC38-Cox8a-GFP cells *in vitro* for 96 hours acquired the GFP signal (Fig. 1E-F), confirming that this transfer occurs in a controlled microenvironment.

We next determined whether the acquired signal represented intact mitochondria or merely degraded protein fragments. We leveraged a single-nucleotide polymorphism in the mitochondrial *mt-Co3* gene between C57BL/6-derived MC38 cells and BALB/c mice^21^. The BALB/c *mt-Co3* sequence contains an G9348A mutation that abolishes AspI restriction site, which is present in the C57BL/6 *mt-Co3* sequence. Following subcutaneous inoculation of MC38 cells into BALB/c mice, we sorted tumor-infiltrating CD8^+^GFP^+^ and CD8^⁺^GFP^-^ T cells, extracted their total DNA, and amplified the *mt-Co3* region. AspI digestion revealed that CD8^+^GFP^-^ cells yielded a single 385-bp band identical to the BALB/c control. Conversely, CD8^+^GFP^+^ cells exhibited a mixed digestion profile indicating the presence of both BALB/c and C57BL/6 mtDNA (Fig. 1G). This indicates that CD8^+^GFP^+^ T cells acquire intact mitochondria containing the donor mitochondrial genome. Parallel imaging flow cytometry revealed that the acquired GFP^+^ mitochondria localized both intracellularly and at the cell surface of CD8^+^ T cells (Fig. 1H-I). Importantly, these acquired GFP^+^ mitochondria could be labelled by Mitotracker Red, confirming that the CD8^+^ T cells had acquired functional mitochondria possessing membrane potential from tumor (Fig. 1H-I).

Finally, we evaluated the organelle specificity of this transfer to rule out trogocytosis or non-specific membrane fusion. We engineered MC38 cell lines co-expressing Cox8a-GFP alongside either a plasma membrane marker (GPR179-mCherry) or an endoplasmic reticulum marker (RCN1-BFP) (Fig. S1H). Tumor-infiltrating CD8^+^ T cells selectively acquired the mitochondrial GFP signal without acquiring the co-expressed membrane or ER markers (Fig. 1J-K). We observe that this intercellular transfer is highly specific to mitochondria and distinct from generic membrane nibbling.

### Mitochondria Transfer is TCR-Independent and Mediated by Cell Contact

Mitochondria can be transferred to recipient cells via contact-independent mechanisms—such as through extracellular vesicles (EVs), exosomes, or “naked” mitochondria directly released into the extracellular space—or via contact-dependent mechanisms, such as tunneling nanotubes (TNTs)^22,23^. To distinguish whether mitochondrial transfer from tumor cells to CD8^+^ T cells requires direct cellular contact, we employed an *in vitro* transwell system that permits the diffusion of soluble factors and vesicles but prevents direct cellular contact. Under these conditions, CD8^+^ T cells failed to acquire mitochondria from the tumor cells (Fig. 2A-B), indicating a strict requirement for cell-to-cell contact for mitochondria to be transferred from tumor cells to CD8^+^ T cells. This contact-dependent mechanism logically implicated TNTs, which are known to mediate organelle transport^24^. Since the Trak1–Miro1 complex is essential for microtubule-based mitochondrial transport via TNTs^25^, we hypothesized that Trak1–Miro1 complex mediates mitochondria transfer to CD8^+^ T cells. Indeed, shRNA-mediated knockdown of either Trak1 or Miro1 in MC38 cells significantly impaired mitochondria transfer from tumor to tumor-infiltrating CD8^+^ T cells (Fig. 2C-D, Fig. S2), supporting a TNT-dependent manner. Corroborating this physical requirement in vivo, confocal microscopy of tumor tissues revealed direct structural interactions between tumor cells and infiltrating CD8^+^ T cells (Fig. 2E). Collectively, these data demonstrate that mitochondrial transfer require direct cell-cell contact and mediated by TNTs in a Trak1–Miro1–dependent manner.

**Fig. 2.**
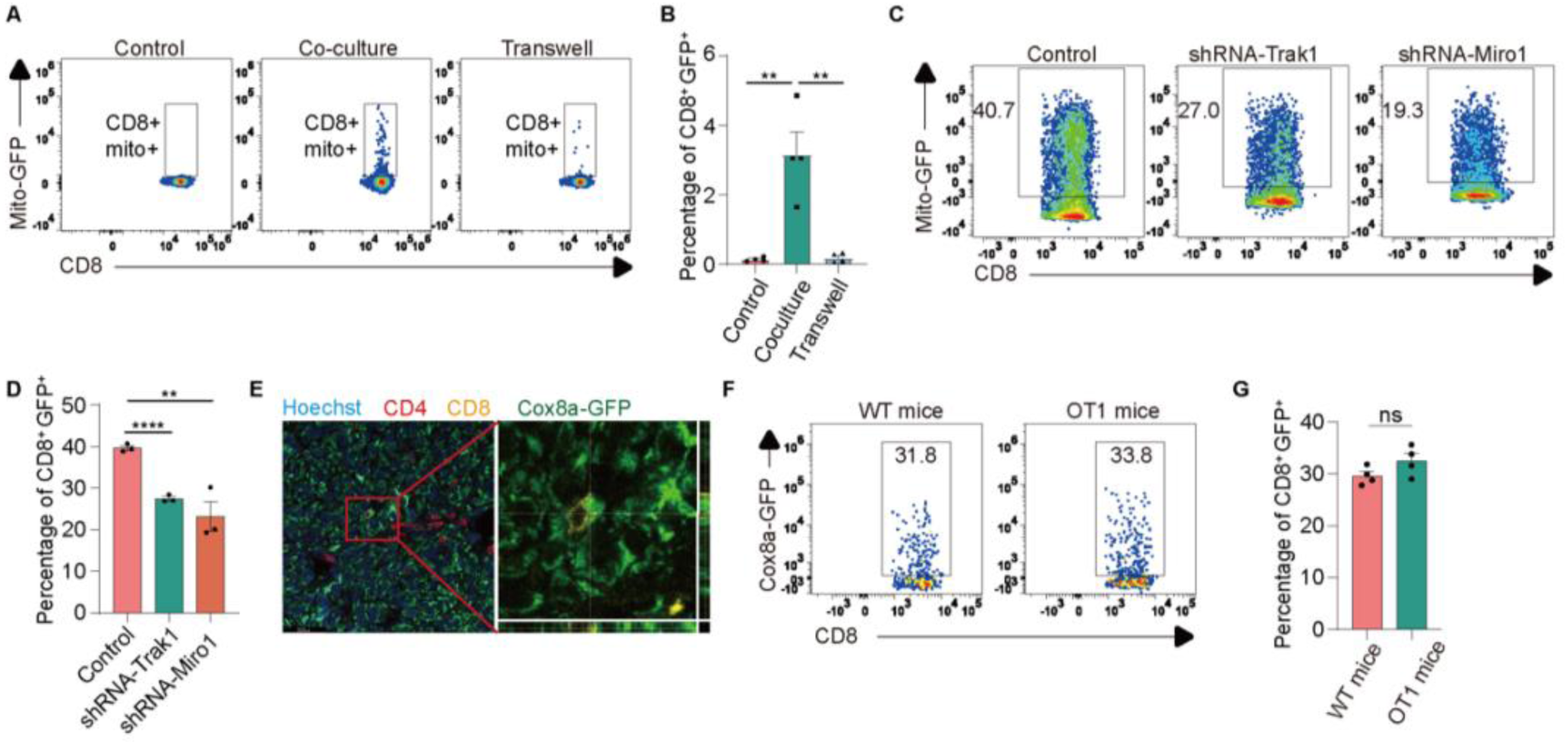
Mitochondria transfer between tumor and T cell depends on cell-cell contact. **A-B**, Analysis of CD8⁺ T cell acquisition of MC38 tumor-derived mitochondria in a transwell co-culture system, as shown in the flow cytometry plot (**A**) and statistical analysis (**B**). **C**–**D**, CD8⁺ T cell acquisition of mitochondria from shRNA knockdown MC38 tumors, as shown in the plot (**C**) and quantitative analysis (**D**). **E**, IHC image depicting a tumor-infiltrating CD8⁺ T cell in contact with a tumor cell. **F-G**, Flow cytometric analysis of *in vivo* mitochondrial transfer from MC38 tumors into CD8⁺ T cells in OT1 mice, shown in a representative plot (**F**) and quantified in (**G**). None Significance: ns (*P* > 0.05), \**P* < 0.05, \*\**P* < 0.01, \*\*\**P* < 0.001, \*\*\*\**P* < 0.0001 (two-way ANOVA and unpaired Student’s t-test).

Because CD8^+^ T cells conventionally engage target cells through T cell receptor (TCR) recognition of cognate antigens^26^, we assessed whether antigen specificity dictates this contact-dependent transfer. We subcutaneously inoculated MC38-Cox8a-GFP cells into OT-I transgenic mice, whose CD8⁺ T cells do not recognize MC38 endogenous antigens. Despite the absence of specific TCR engagement, OT-I CD8⁺ T cells acquired tumor mitochondria at frequencies comparable to wild-type controls (Fig. 2G).

Collectively, these data establish that tumor-to-T cell mitochondrial transfer is a contact-dependent, TCR-independent process mediated by the Trak1–Miro1 transport machinery.

### Mitochondrial transfer enhances CD8 cell cytotoxicity

To determine how tumor-derived mitochondria impact CD8⁺ T cell function, we profiled canonical activation and exhaustion signatures. We observed no significant differences in the expression of activation (CD44, CD62L) or exhaustion (PD-1, TIM-3) markers between CD8^+^GFP^+^ and CD8^+^GFP^-^ populations (Fig. S3A-D), indicating that mitochondrial acquisition does not globally alter these standard differentiation states. However, upon ex vivo pharmacological stimulation with PMA, tumor-infiltrating CD8^+^GFP^+^ cells produced significantly higher protein levels of the cytotoxic effector molecules Granzyme B and Perforin compared to their GFP^-^counterparts (Fig. 3A-C).

**Fig. 3.**
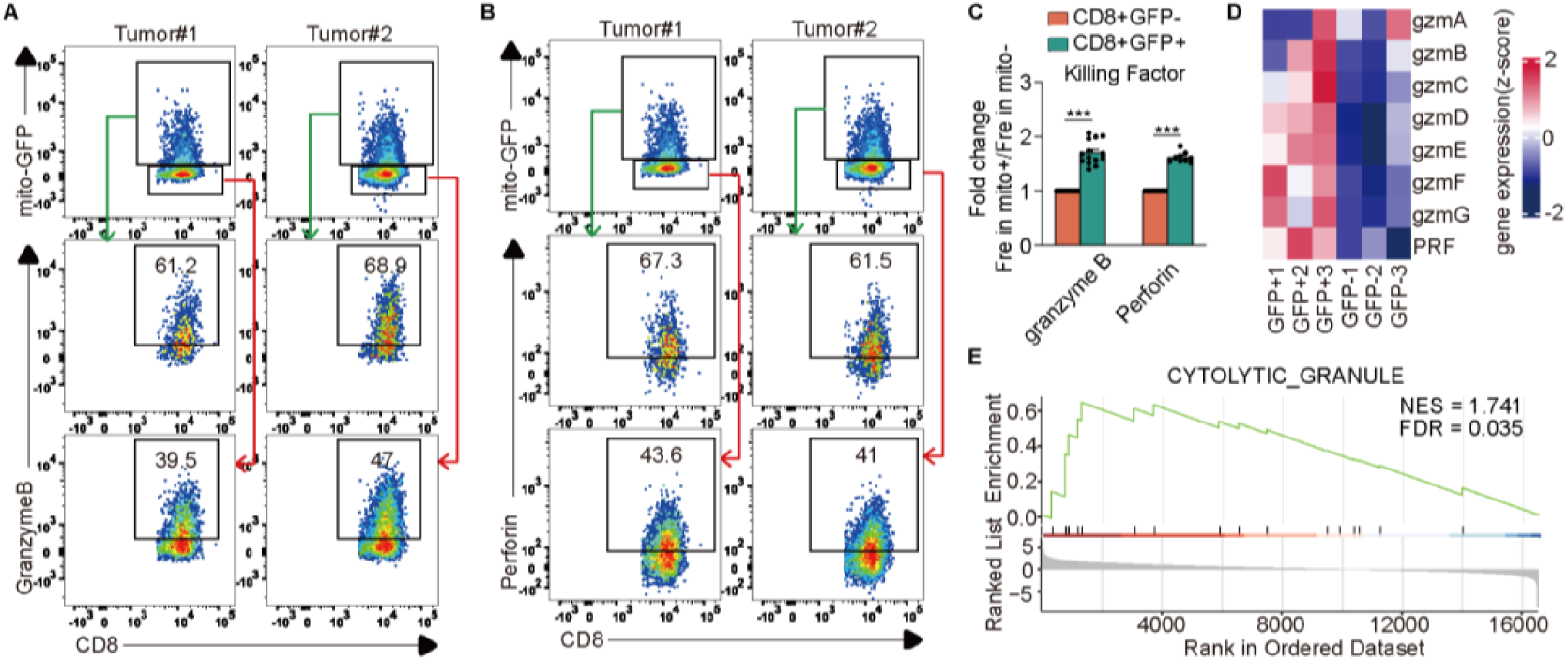
CD8⁺ mito⁺ T cells exhibit enhanced production of antitumor effector molecules. **A-C**, Expression of granzyme B (GzmB) and perforin in tumor-infiltrating CD8⁺ T cells following *ex vivo* PMA/ionomycin stimulation, presented as a graphical plot (**A, B**) and quantitative analysis (**C**). **D**, RNA-seq profiling of granzyme family members and perforin expression in tumor-infiltrating CD8⁺ T cells. **E**, GSEA analysis of CYTOLYTIC_GRANULE pathway. None Significance: ns (*P* > 0.05), \**P* < 0.05, \*\**P* < 0.01, \*\*\**P* < 0.001, \*\*\*\**P* < 0.0001 (unpaired Student’s t-test).

To map the transcriptional basis underlying this functional difference, we performed RNA-sequencing on sorted tumor-infiltrating CD8^+^GFP^+^ and CD8^+^GFP^-^ populations. Transcriptomic analysis revealed a broad upregulation of genes encoding cytotoxic effector molecules, including Prf1 and multiple granzymes (Gzmb, Gzmc, Gzmd, Gzme, Gzmf, Gzmg), in CD8^+^GFP^+^ cells (Fig. 3D). This transcriptional profile strictly aligns with the elevated effector protein expression observed post-stimulation (Fig. 3C). Concordantly, Gene Set Enrichment Analysis (GSEA) demonstrated a significant enrichment of cytotoxic granule-related pathways within the CD8^+^GFP^+^ population (Fig. 3E).

Collectively, these data demonstrate that the acquisition of tumor mitochondria selectively bolsters the cytotoxic transcriptional program and effector capacity of CD8^+^ T cells, independent of their global activation or exhaustion status.

### S1P-S1PR1 mediates the enhanced cytotoxic molecules secretion in CD8^+^GFP^+^ T cells

To delineate the mechanism driving enhanced cytotoxicity in CD8^+^GFP^+^ T cells, we analyzed our RNA-seq data and found that CD8^+^GFP^+^ cells exhibited significantly lower mRNA expression of *S1pr1* (Fig. 4A), which encodes the membrane-bound receptor for sphingosine-1-phosphate (S1P). Because the transcriptional downregulation of *S1pr1* is known to maintain CD8^+^ T cell effector function^27^, we hypothesized that this reduction contributes to the potentiated cytotoxic phenotype observed in CD8^+^GFP^+^ cells. To validate this in vivo, we reanalyzed publicly available scRNA-seq datasets from both B16^28–32^ and MC38^32–35^ tumor models. Within the CD8^+^ T cell clusters of both models, lower *S1pr1* expression consistently correlated with elevated levels of *Prf1* and *Gzmb* (Fig. 4B–C). These findings confirm a robust inverse relationship between *S1pr1* transcription and cytotoxic effector capacity.

**Fig. 4.**
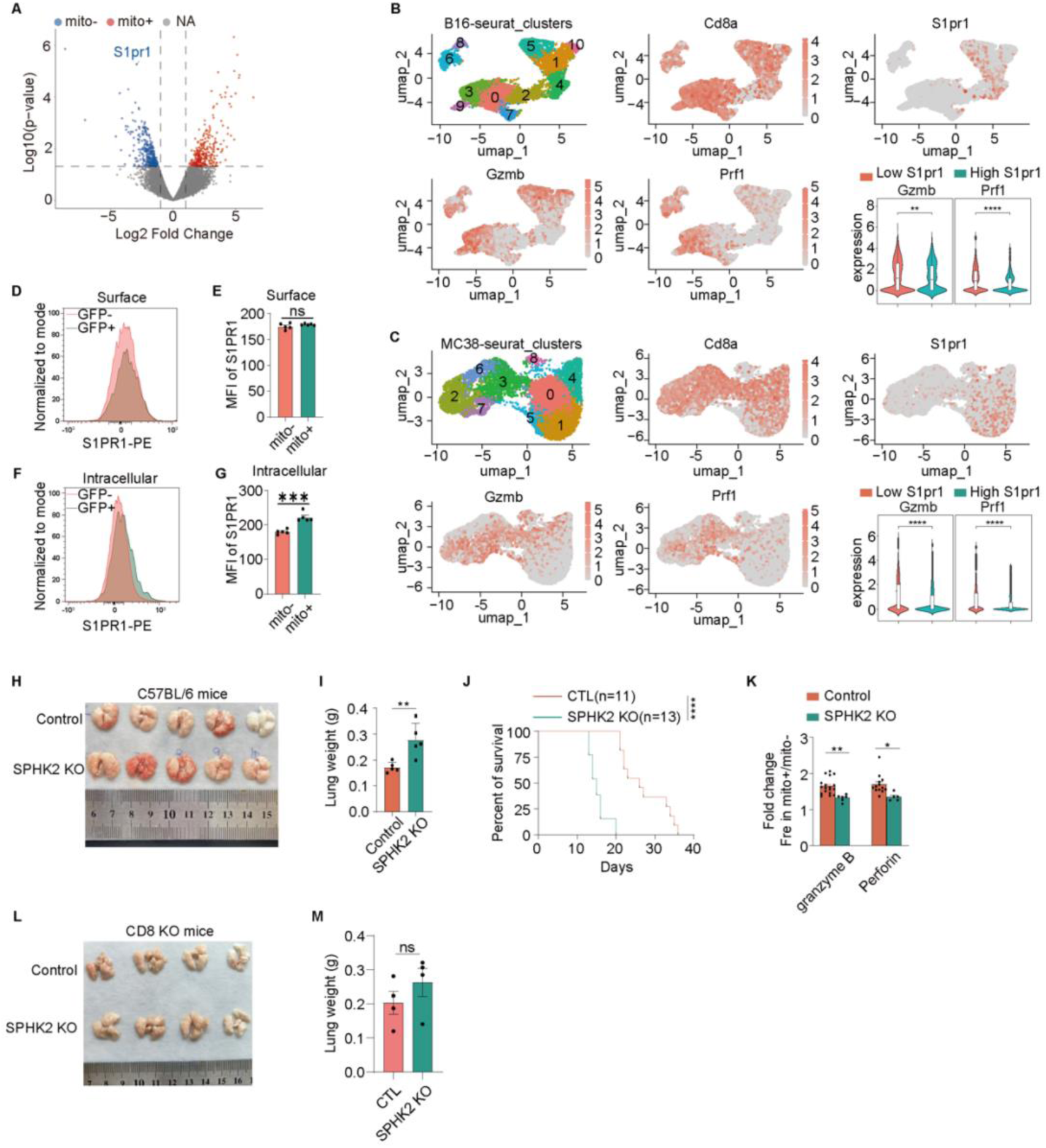
S1P-S1PR1 mediates the enhanced cytotoxic molecules secretion in CD8^+^ mito^+^ T cells. **A**, Volcano plot of gene expression in CD8⁺ mito⁺ versus CD8⁺mito⁻ T cells. **B-C**, Expression profiles of S1PR1, granzyme B, and perforin in tumor-infiltrating CD8⁺ T cells, visualized by UMAP. Data are derived from published datasets for B16 (**B**) and MC38 (**C**) models. **D-G**, Comparative surface (**D, E**) and intracellular (**F, G**) S1PR1 receptor levels in CD8⁺ mito⁺ versus mito⁻ T cells. **H**-**J**, Lung gross morphology (**H**), weight (**I**), and survival (**J**) of mice inoculated with SPHK2 KO MC38 cells. **K**, Analysis of granzyme B and perforin in *ex vivo*-stimulated tumor-infiltrating CD8⁺ T cells from SPHK2-KO MC38 tumor models. **L-M**, Lung gross morphology (**L**) and weight (**M**) of CD8 KO mice inoculated with MC38 cells. None Significance: ns (*P* > 0.05), \**P* < 0.05, \*\**P* < 0.01, \*\*\**P* < 0.001, \*\*\*\**P* < 0.0001 (unpaired Student’s t-test). Survival curves were compared using the log-rank test, and statistical significance was defined as \*\*\*\**P* < 0.0001.

Because S1P–S1PR1 signaling regulates the production of Perforin and Granzyme B in CD8^+^ T cells^15^, and that S1P generated by SPHK2 can localize to mitochondria^10^, we postulated that tumor-derived mitochondrial S1P activates S1PR1 on recipient CD8^+^GFP^+^ cells. Ligand-induced activation of S1PR1 triggers its rapid internalization^36^. Consistent with this dynamic, CD8^+^GFP^+^ cells exhibited elevated intracellular S1pr1 protein levels without influencing its surface expression level (Fig. 4D-G), indicative of active ligand-receptor engagement and subsequent internalization. To determine whether tumor-derived S1p drives this cytotoxic enhancement, we generated *Sphk2*-knockout (*Sphk2*-KO) MC38 cells (Fig. S4A). While *Sphk2* deficiency did not alter tumor cell proliferation *in vitro* (Fig. S4B), it accelerated subcutaneous tumor growth (Fig. S4C-D), exacerbated pulmonary metastatic burden (Fig. 4H-I), and reduced the overall survival of tumor-bearing mice (Fig. 4J). Concomitantly, CD8^+^GFP^+^ T cells isolated from *Sphk2*-KO tumors exhibited significantly reduced expression of Granzyme B and Perforin (Fig. 4K). This indicates that tumor-derived S1p—likely transferred via the acquired mitochondria—is required to potentiate CD8^+^ T cell cytotoxicity. Finally, the exacerbated metastatic phenotype observed in C57BL/6 mice bearing *Sphk2*-KO tumors was completely abrogated in CD8^+^ T cell-deficient hosts (Fig. 4L-M), confirming that this effect is strictly dependent on CD8^+^ T cell function.

Collectively, these findings support a model in which Sphk2-dependent, tumor-derived S1P—likely delivered via intercellular mitochondrial transfer—activates S1PR1 on recipient CD8^+^ T cells, driving receptor internalization and potentiating anti-tumor cytotoxicity. Collectively, these findings support a model wherein tumor-derived mitochondria, potentially via SPHK2-generated mitochondrial S1P, activate S1PR1 on CD8⁺ T cells, leading to receptor internalization and potentiation of cytotoxic function.

## Discussion

The tumor microenvironment (TME) harbors complex intercellular communication networks that critically regulate anti-tumor immunity, yet the role of organelle transfer in shaping T cell function remains poorly defined. Our study reveals a novel intercellular communication pathway within the tumor microenvironment: functional mitochondria can be transferred from tumor cells to infiltrating CD8⁺ T cells in a TCR-independent, cell-contact-dependent manner. Remarkably, this transfer does not induce canonical T cell activation or exhaustion but instead specifically potentiates the cytotoxic machinery of recipient CD8⁺ T cells. Mechanistically, we demonstrate that this functional enhancement is mediated by the sphingolipid signaling axis: tumor-derived mitochondria supply SPHK2-generated mitochondrial S1P, which activates S1PR1 on recipient CD8⁺ T cells and induces receptor internalization—ultimately boosting the cytotoxic effector program of these cells. These findings position tumor-to-CD8⁺ T cell mitochondrial transfer as a previously underappreciated modulator of anti-tumor immunity, challenging the conventional view of tumor cells as solely immunosuppressive and uncovering a novel pro-immunogenic mechanism within the TME.

The transfer of mitochondria from cancer cells to immune cells has been sporadically reported, often in the context of dysfunction or metabolic reprogramming^7,37^. The selectivity for mitochondrial transfer—over other organelles—and its occurrence via structures reminiscent of TNTs suggest an active, regulated process rather than passive debris uptake. This observation aligns with the emerging role of TNTs in facilitating the exchange of functional organelles in stress responses^38^. Critically, our finding that tumor-to-CD8⁺ T cell mitochondrial transfer enhances, rather than suppresses, cytotoxic function directly challenges the prevailing view that tumor cells primarily subvert immune responses through intercellular exchange. This novel observation reveals that the TME harbors cryptic pro-immunogenic signals—specifically, tumor-derived functional mitochondria and their associated S1P-S1PR1 signaling—that can be harnessed to boost anti-tumor immunity.

Among our findings, the most intriguing is the link between acquired mitochondria and the S1P-S1PR1 pathway. S1P is a pleiotropic lipid mediator, and its role in lymphocyte egress from lymphoid organs via S1PR1 is well-established^13^. However, its function within solid tumors, particularly in regulating T cell effector function, is less clear. Our data, showing S1PR1 internalization specifically in CD8⁺GFP⁺ cells and the dependency of the phenotype on tumor SPHK2, propose a model where tumor mitochondria serve as a vehicle for localized S1P delivery. This localized delivery of S1P—directly from tumor mitochondria to CD8^+^ T cells—enables precise activation of S1PR1 at the tumor-T cell interface, thereby bypassing the inhibitory effects of high systemic or TME-resident S1P that often constrain T cell function^39,40^. However, we acknowledge that definitive proof requiring direct demonstration of mitochondrial S1P activating CD8^+^ T cell S1PR1, for instance via genetic rescue with mitochondrially-targeted SPHK2, is an important future direction.

Our study also raises several critical unanswered questions that warrant further investigation, which will help to fully dissect the biological significance of tumor-to-CD8⁺ T cell mitochondrial transfer. First, what molecular mechanisms govern the selectivity of mitochondrial transfer to CD8^+^ T cells (and potentially specific subsets thereof)? Molecules present on the tumor mitochondrial surface, or the activation and metabolic states of recipient CD8⁺ T cells, may serve as key determinants of this selective transfer, and their identities deserve detailed exploration. Second, while our study focused on the potentiation of CD8⁺ T cell cytotoxicity, the acquisition of intact, functional tumor mitochondria—adapted to the hypoxic and nutrient-depleted tumor microenvironment (TME)—is likely to profoundly reshape the metabolic landscape of recipient CD8⁺ T cells. These tumor-adapted mitochondria may supply critical metabolic intermediates to fuel biosynthetic pathways and sustain high-efficiency effector function, thereby overcoming the metabolic constraints that often drive T cell exhaustion in the TME. Future metabolomic profiling and Seahorse extracellular flux analyses of sorted CD8⁺GFP⁺ and CD8⁺GFP⁻ T cell populations will clarify the specific metabolic reprogramming induced by tumor mitochondrial transfer. Third, the clinical translational potential of the tumor-to-CD8⁺ T cell mitochondrial transfer pathway remains to be fully explored, which is critical for translating our basic findings into novel cancer immunotherapeutic strategies.

In conclusion, our study identifies tumor-to-CD8⁺ T cell mitochondria transfer as a novel, cell-contact-dependent, and TCR-independent intercellular communication mechanism that selectively potentiates CD8⁺ T cell cytotoxicity in the tumor microenvironment. Furthermore, our findings reveal that the SPHK2-derived mitochondrial S1P-S1PR1 signaling axis mediates this cytotoxic potentiation, offering a novel perspective on how lipid metabolism and organelle intercellular transfer intersect to regulate anti-tumor immunity—challenging the prevailing view that tumor cells primarily subvert immune function through intercellular communication. Collectively, this tumor-to-CD8⁺ T cell mitochondrial transfer pathway—together with its downstream SPHK2-S1P-S1PR1 regulatory axis—represents a promising therapeutically targetable node to enhance CD8⁺ T cell-mediated anti-tumor immunity, with potential applications in improving the efficacy of adoptive T cell therapy and other cancer immunotherapies. While our study establishes the core mechanisms and functional consequences of this transfer, future investigations addressing the unresolved questions outlined above will further advance our understanding of this novel intercellular communication mode and its translational potential.

## Methods

### Key resources table

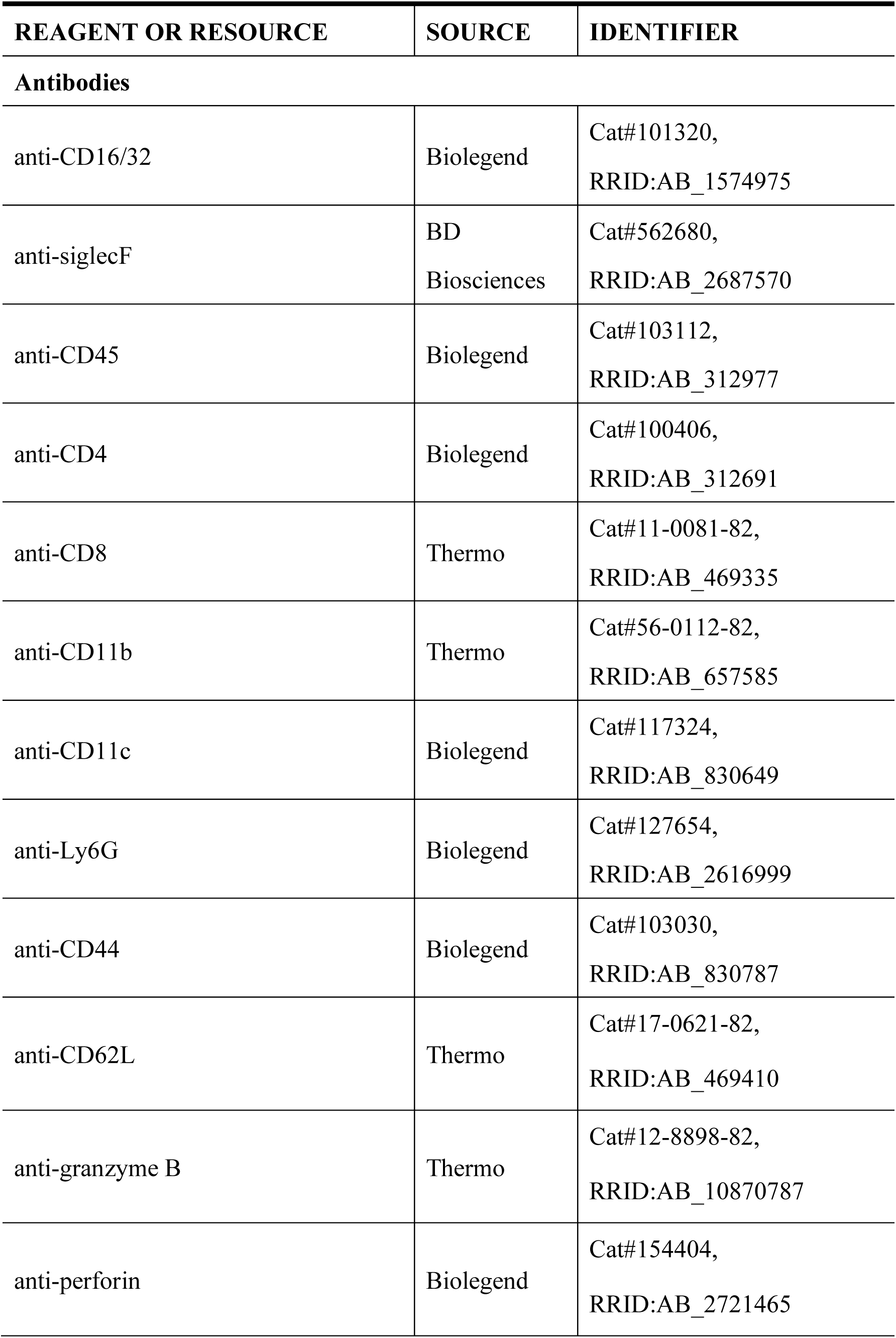
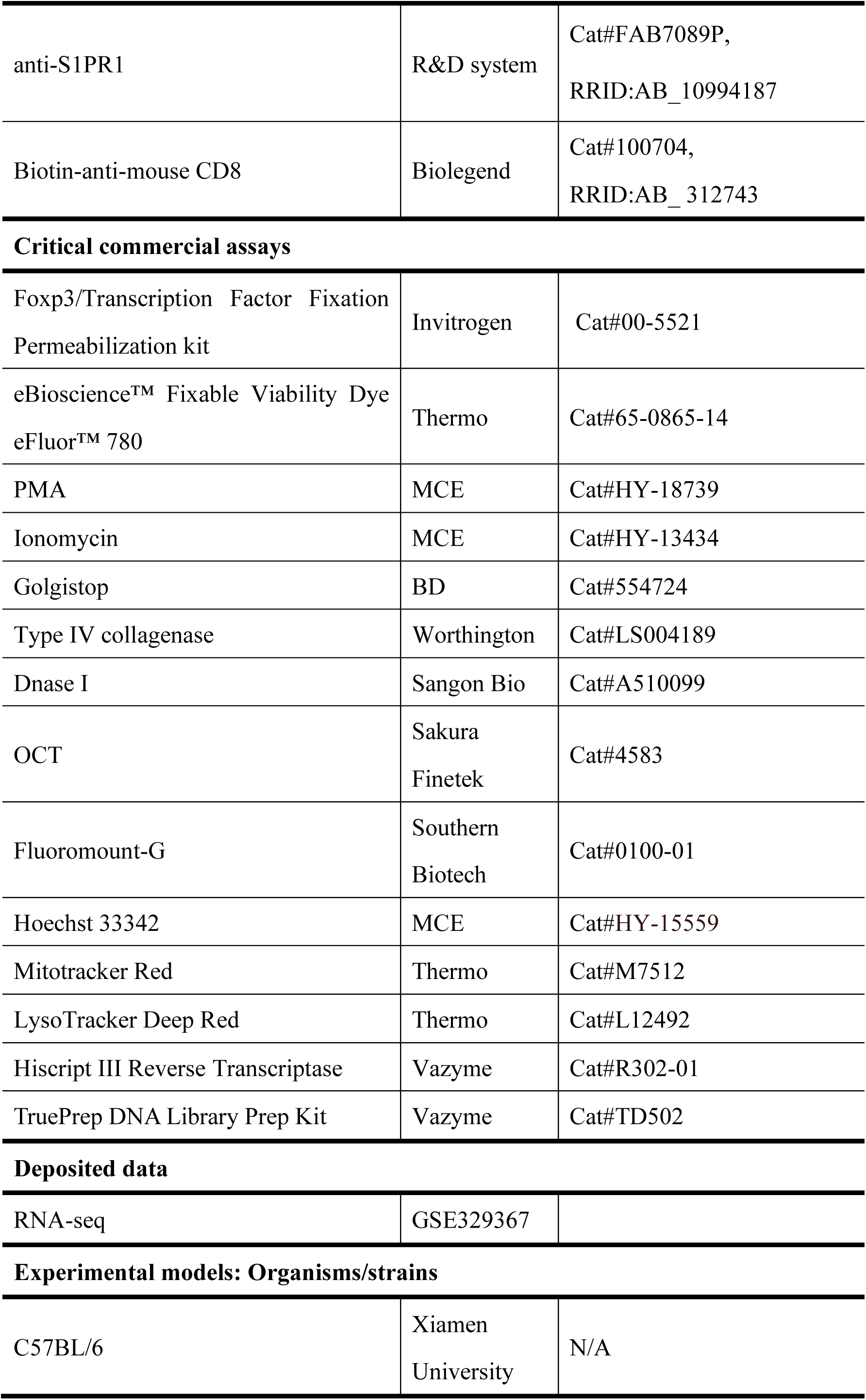
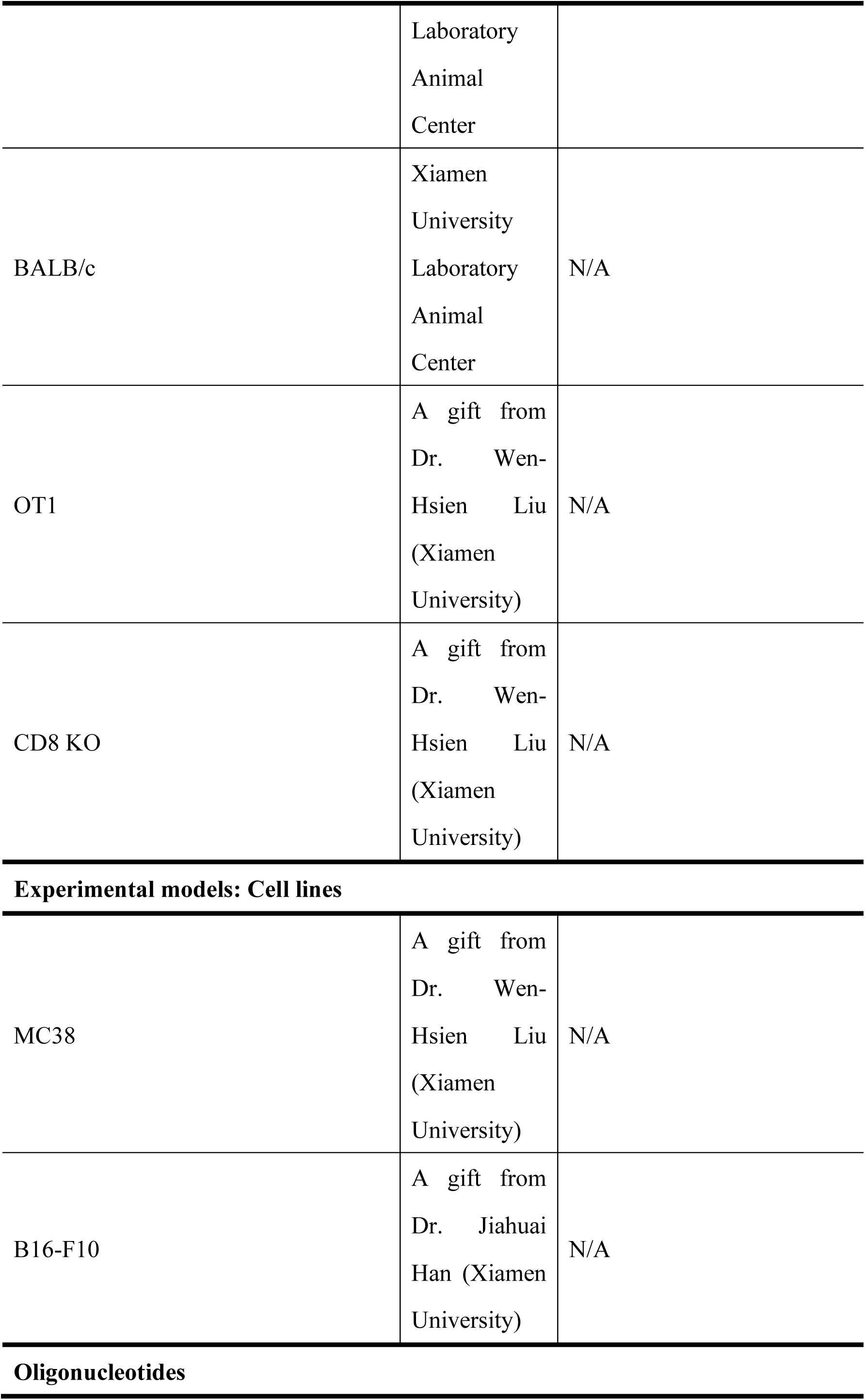
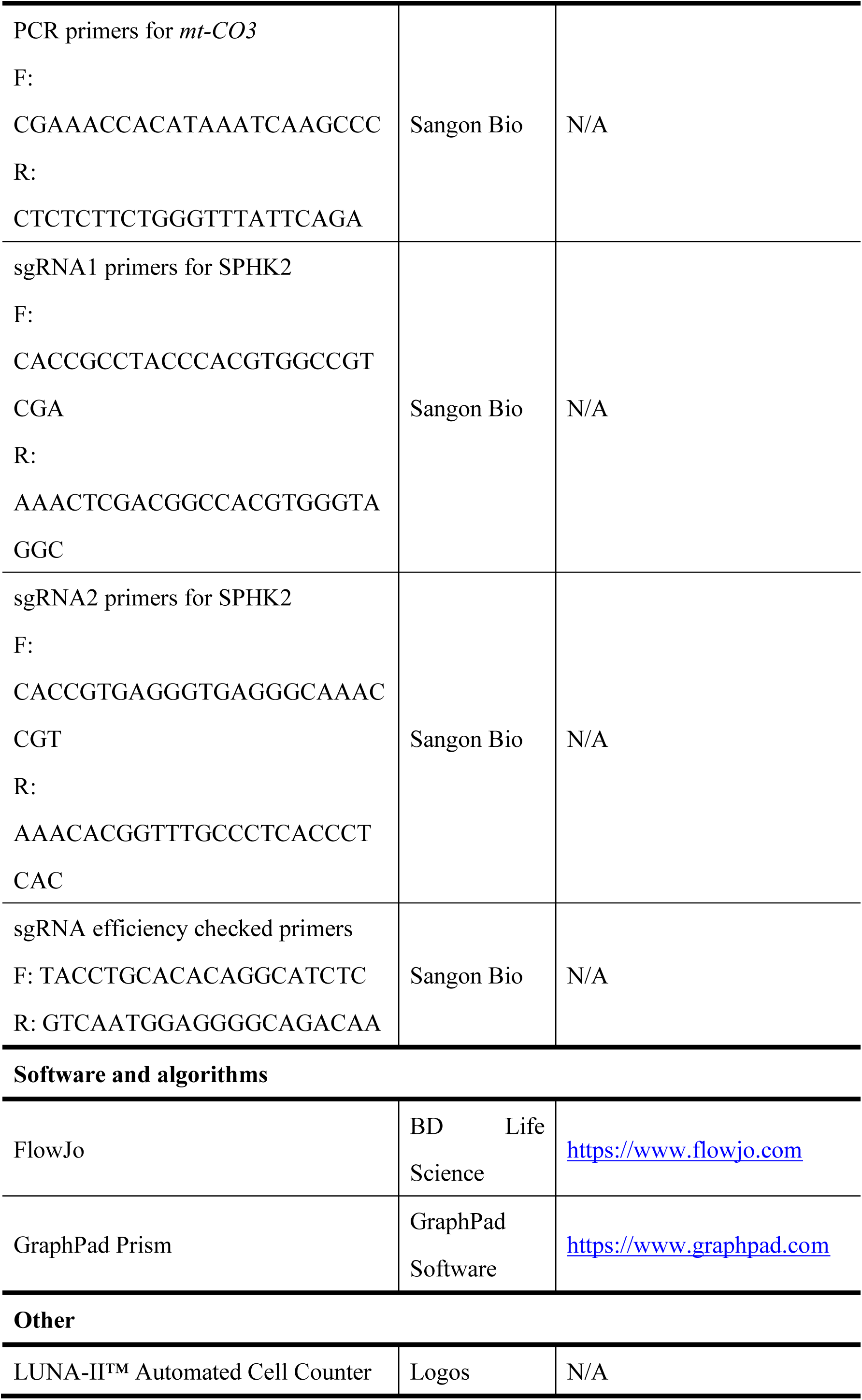
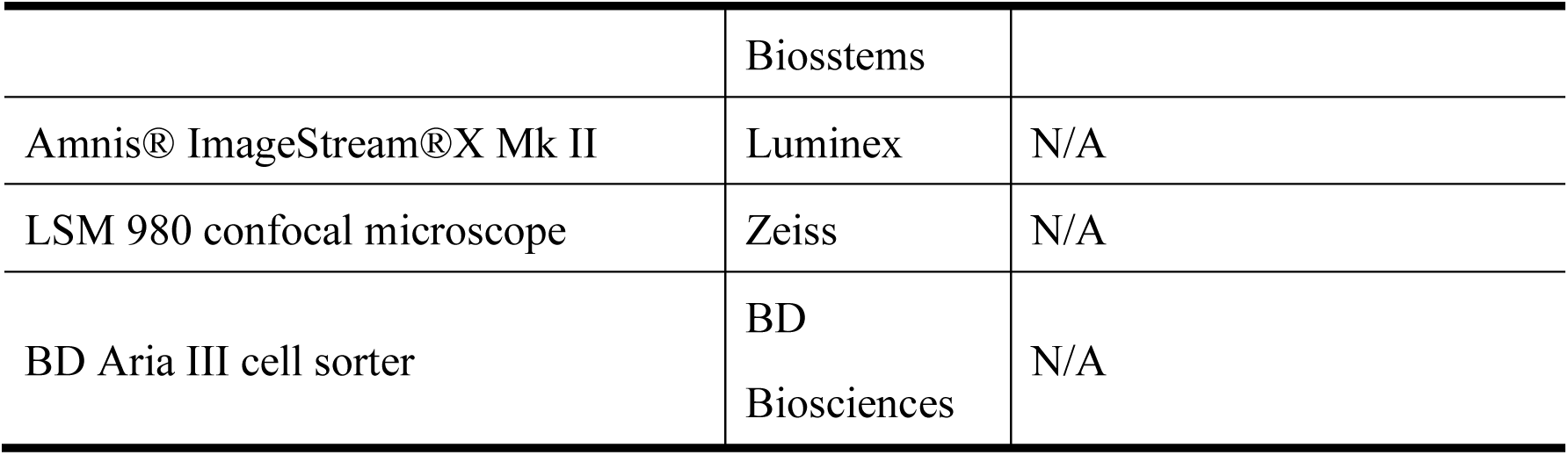

### Mice

Wild-type C57BL/6 and BALB/c mice were sourced from the Laboratory Animal Center of Xiamen University. OT-I and CD8 KO mice were kindly gift by Dr. Wen-Hsien Liu (Xiamen University). 8–12-week-old, sex-matched mice were used for experiments. All mice were maintained in specific pathogen–free (SPF) barrier facilities under standardized conditions. Animal experiments were approved by the Xiamen University Institutional Animal Care and Use Committee and conducted in accordance with institutional ethical guidelines.

### Cell lines

Cell Line: MC38; Species/Origin: mouse, immortalized colon cancer cell line; Maintenance and Care: Cells were cultured in Dulbecco’s Modified Eagle Medium (DMEM) supplemented with 10% (v/v) fetal bovine serum (FBS) and 1% penicillin-streptomycin. Cell Line: B16-F10; Species/Origin: mouse, immortalized melanoma cell line; Maintenance and Care: Cells were cultured in Dulbecco’s Modified Eagle Medium (DMEM) supplemented with 10% (v/v) fetal bovine serum (FBS) and 1% penicillin-streptomycin. All cells were cultured at 37℃ in a humidified atmosphere of 5% CO^2^. Cells were routinely tested for mycoplasma contamination using Mycoplasma qPCR Detection Kit (C0303S, Beyotime Biotechnology, Shanghai, China)

## Method details

### *In vivo* tumor models

B16 and MC38 tumor cells tagged by mitochondrial located fluorescent protein were subcutaneously inoculated or intravenous injection into wildtype C57BL/6 mice (1 × 10^6^ MC38 or 2.5 × 10^5^ B16-F10 tumor cells per mouse). For TCR dependent assay, MC38-Cox8a-GFP tumor cells were subcutaneously inoculated into OT-I mice. For single nucleotide polymorphism enzyme digestion experiments, MC38-Cox8a-GFP tumor cells were subcutaneously inoculated into BALB/c mice. For CD8 dependent assay, SPHK2 KO MC38-Cox8a-GFP tumor cell were administered via tail vein injection into CD8 KO mice. All mice were house in specific pathogen-free conditions at Xiamen University Laboratory Animal Center. Institutional Animal Care and Use Committee approved all mouse experiments defined by the Xiamen University Laboratory Animal Center. We have complied with all relevant ethical regulations for animal use.

### Preparation of Single-Cell Suspension

Freshly excised tumors were immediately placed on ice and processed into single-cell suspensions. In brief, tumor tissues were cut into pieces and digested with 1 mg/mL type IV collagenase, 50 U/mL DNase I in DMEM medium at 37 °C for 20 min in a shaking incubator. The suspension was passed through a 70-µm cell strainer, and then red blood cells were lysed with ACK to prepare a single-cell suspension.

### Flow cytometry analysis

Following single-cell suspension preparation, 3 × 10^6^ cells were blocked with anti-CD16/32 antibody, and then cells were incubated with specific cell surface marker. For coculture assay, co-cultured cells were harvest for intracellular staining,

### *In vitro* co-culture assay

CD8⁺ T cells from splenocytes were collected with BD FACSAria III cell sorter, stimulated with CD3 (3 μg/mL) and CD28 (3 μg/mL) antibody for 12 h, and then co-cultured with MC38-Cox8a-GFP cells for 96h. For transwell assay, CD8^+^ T cells were seeded in the bottom layer, while tumor cells were seeded in the upper layer. The transwell has a pore size of 0.4 μm.

### Constructs, lentivirus production and transfection

Cox8a-GFP, Cox8a-BFP and Tomm20-GFP were clone into lentivirus backbone plasmid (PLV) for further transfection. For Cox8a-GFP and Cox8a-BFP, Cox8a located sequence were fused with GFP or BFP. For Tomm20-GFP, PCR amplified whole Tomm20 CDS region were fused with GFP. For lentivirus production, PLV vectors encoding target sequence were co-transfected with packaging plasmids (PMD2G and PSPAX) into 293T cells. After 48h, the supernatants were harvest and transduced into MC38 and B16 tumor cells.

### Imaging Flow Cytometry

Single-cell tumor suspensions were analyzed by imaging flow cytometry (Amnis ImageStream MK II) following staining with Hoechst 33342, a fixable viability dye, anti-CD8 antibody, and MitoTracker Red.

### Immunofluorescence Staining

Tumor tissues were dehydrated in 30% sucrose for 16–20 h, and embedded in OCT compound. Cryosections (25 μm) were prepared using a CM1905 cryostat (Leica). Sections were rehydrated with PBS and blocked with a solution containing 1% normal serum (mouse, rat, and rabbit), 1% bovine serum albumin (BSA), and 0.2% Triton X-100 in PBS. Subsequently, sections were incubated directly with a mixture of fluorophore-conjugated primary antibodies, including anti-mouse CD8a and anti-mouse CD4 for 24 h at room temperature in a dark humidified chamber. The sections were then stained with Hoechst 33342 solution for 10 min at room temperature, followed by five 10-min washes with PBS and coverslipped with Fluoromount-G.

### Fluorescence microscopy

To confirm mitochondrial localization of the expressed fluorescent proteins, B16 and MC38 cells were co-stained with MitoTracker Red. Images were acquired using a Zeiss LSM 980 confocal microscope. For imaging of tumor-infiltrating macrophages, cells were first sorted using a BD FACS Aria III, then stained with LysoTracker and MitoTracker, and finally imaged on a Zeiss LSM 980 confocal microscope. For tumor section, stained sections were imaged using a Leica TCS SP8 confocal microscope.

### RNA-seq Library Construction

MC38 tumor-infiltrating CD8^+^mito^+^ and CD8^+^mito^-^ population were sorted with BD ARIA III cell sorter, and RNA was extracted and reverse transcribed according to the Hiscript III Reverse Transcriptase kit following the manufacturer’s instructions. Sequencing libraries were generated using TruePrep DNA Library Prep Kit V2 for Illumina (Vazyme), and index codes were added to attribute sequences to each sample. The library was sequenced on an Illumina Novaseq platform, and 150 bp paired-end reads were generated.

### RNA-seq data analysis

RNA-seq paired-end reads were aligned to the mouse reference genome (GRCm38) using HISAT2 with default parameters. Prior to alignment, adapter sequences and low-quality bases were trimmed. Read counts per gene were generated using featureCounts, and FPKM values were calculated based on gene length and these counts. Differential expression analysis was performed with DESeq2, comparing two biological replicates. Genes with a fold change >2 and an adjusted *p*-value <0.05 were considered significantly differentially expressed. Pathway enrichment analysis was conducted using ClusterProfiler (R) with a significance cutoff of *p*-value < 0.05. All heatmaps were generated using the ComplexHeatmap R package.

### Integration and Re-analysis of Multi-source Single-cell RNA Sequencing Datasets

Public single-cell RNA sequencing (scRNA-seq) datasets for B16 (GSE236680, GSE247640, GSE261553, GSE227699, GSE217038) and MC38 (GSE263524, GSE197229, GSE264446, GSE217038) tumors were downloaded and re-analyzed to interrogate the expression of target genes.

### Generation of SPHK2-knockout (SPHK2) MC38 cell line

The SPHK2-knockout (SPHK2-KO) MC38 cell line was generated via CRISPR-Cas9-mediated genome editing. Two sgRNAs (sequence were listed in key resources table) were designed to target CDS of the murine *Sphk2* gene (NCBI Reference Sequence: NM_203280.3). Co-infection of these sgRNAs with Cas9 lentivirus was performed to induce double-strand breaks at both sites, leading to the deletion of a 166-bp fragment (corresponding to genomic coordinates Chr7: 45,362,989 – 45,362,823 on the GRCm39/mm39 assembly). Successful deletion was confirmed by genomic PCR across the edited locus.

### *mt-Co3* sequence amplification and AspI restriction

Total DNA was extracted from the sorted CD8^+^GFP^+^ and CD8^+^GFP^-^ T cells, and the *mt-Co3* fragment was amplified using PCR primers (list in key resources table). Subsequently, the fragment was incubated with AspI Endonuclease at 37°C for 60 minutes, and the results were analyzed via gel electrophoresis.

## Quantification and Statistical Analysis

In the present study, all experiments were conducted using at least two biological replicates. All statistical analyses were conducted using GraphPad Prism v9.0.0. Data are presented as mean ± SEM. Two-tailed unpaired Student’s t-test was used to evaluate the comparison between groups. Survival curves were compared using the log-rank test. Statistical significance was defined as \**p* < 0.05, \*\**p* < 0.01, \*\*\**p* < 0.001, and \*\*\*\**p* < 0.0001.

## Data availability

The RNA-seq data generated in this study have been deposited in the NCBI Gene Expression Omnibus (GEO) database and are accessible under accession number GSE329367.

## Author Contributions

Shih-Chin Cheng conceived and designed the study. Chuanfang Chen, Xiunan Wang and Haige Li performed the experiments. Data analysis and interpretation were conducted by Chuanfang Chen, Xiunan Wang, Haige Li, Qingxiang Gao, Jia Zhang and Shih-Chin Cheng. Chuanfang Chen and Xiunan Wang drafted the initial manuscript. Jia Zhang and Shih-Chin Cheng supervised the project.

## Competing interests

The authors declare no competing interests.

## Acknowledgments

We acknowledged Dr. Wen-Hsien Liu (Xiamen University, Fujian Province, China) for providing OT-I and CD8 KO mice kindly. Shih-Chin Cheng gratefully acknowledges financial support from the National Natural Science Foundation of China (grants 32161133020), the Fundamental Research Funds for the Central Universities (grant 20720220003), and the Start-up Fund of Xiamen University.

---

**Fig. S1.**
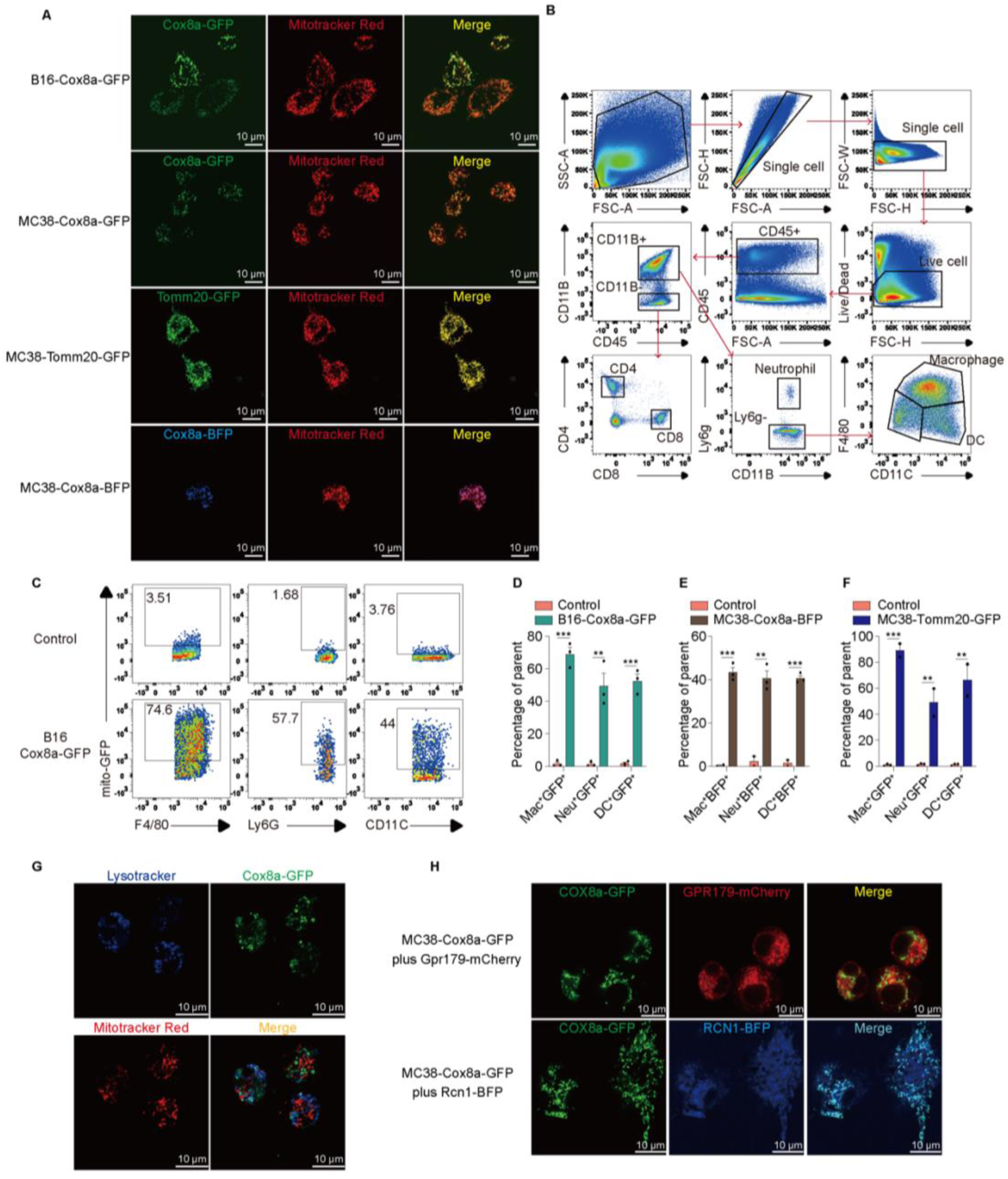
Mitochondrial transfer from tumor cells to immune cells **A**, Representative photographs of MC38 and B16-F10 tumor cell expressing mitochondria targeted fluorescent protein. **B**, Gating strategy of tumor-infiltrated immune cells**. C**-**F**, Flow cytometric analysis of *in vivo* tumor mitochondrial acquisition by macrophages, neutrophils, and DCs, depicted in a representative plot (**C**) and quantified in (**D-F)**. **G**, Representative images of tumor-infiltrating macrophages containing internalized mitochondria. **H**, MC38 cells expressing the indicated fluorescent protein markers, with Cox8a-GFP used as a mitochondrial marker in all cases. (Top) Cells co-expressing GRP179-mCherry. (Bottom) Cells co-expressing RCN1-BFP. None Significance: ns (*P* > 0.05), \**P* < 0.05, \*\**P* < 0.01, \*\*\**P* < 0.001, \*\*\*\**P* < 0.0001 (unpaired Student’s t-test).

**Fig. S2.**
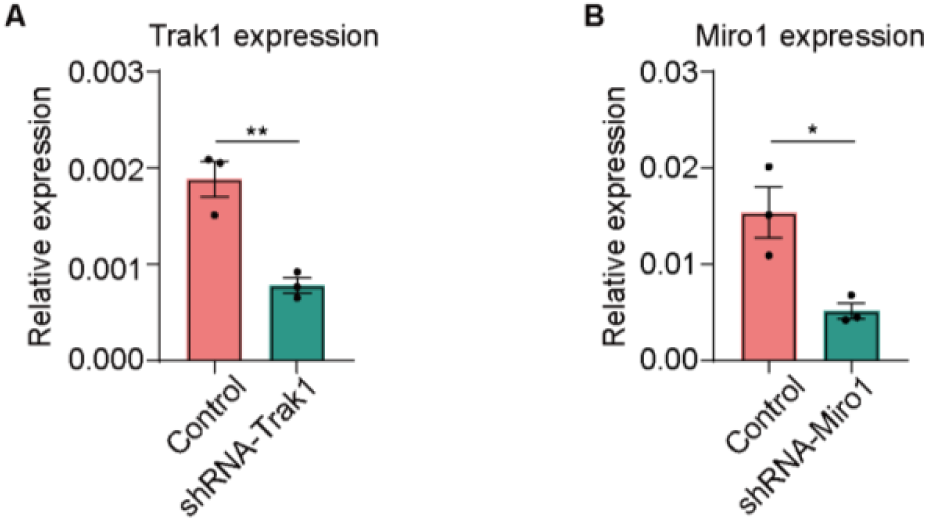
Trak1 and Miro1 shRNA knockdown efficiency. **A-B**, Knockdown efficiency of Trak1 (**A**) and Miro1 (**B**) with shRNA in MC38 cells. None Significance: ns (*P* > 0.05), \**P* < 0.05, \*\**P* < 0.01, \*\*\**P* < 0.001, \*\*\*\**P* < 0.0001 (unpaired Student’s t-test).

**Fig. S3.**
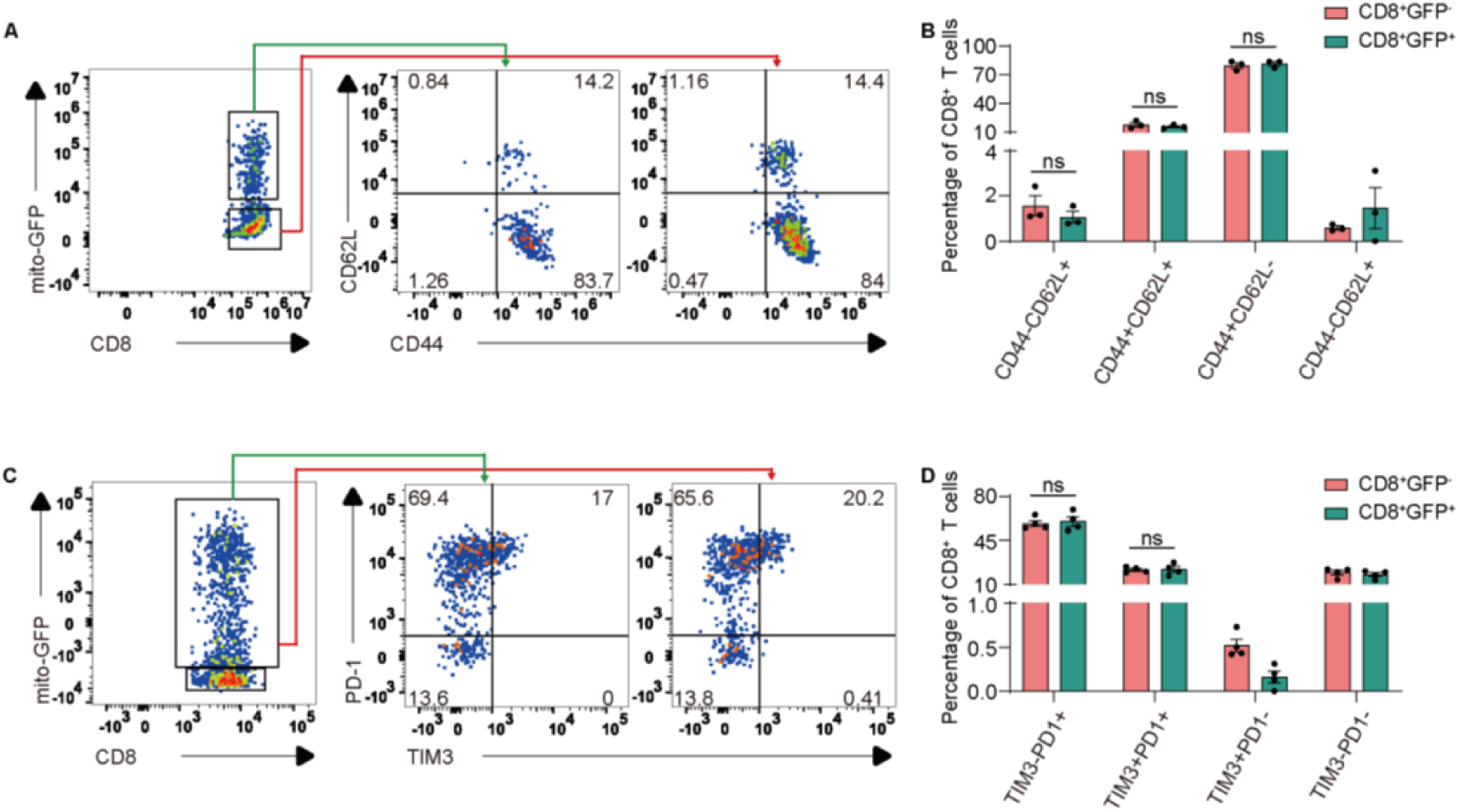
Tumor mitochondrial transfer have no effect on CD8 T cell activation and exhaustion **A-D**, Expression of activation marker (CD44, CD62L) and Exhausted marker (TIM3, PD-1) in tumor-infiltrating CD8⁺ T cells, presented as a graphical plot (**A, C**) and quantitative analysis (**B, D**). None Significance: ns (*P* > 0.05), \**P* < 0.05, \*\**P* < 0.01, \*\*\**P* < 0.001, \*\*\*\**P* < 0.0001 (unpaired Student’s t-test).

**Fig. S4.**
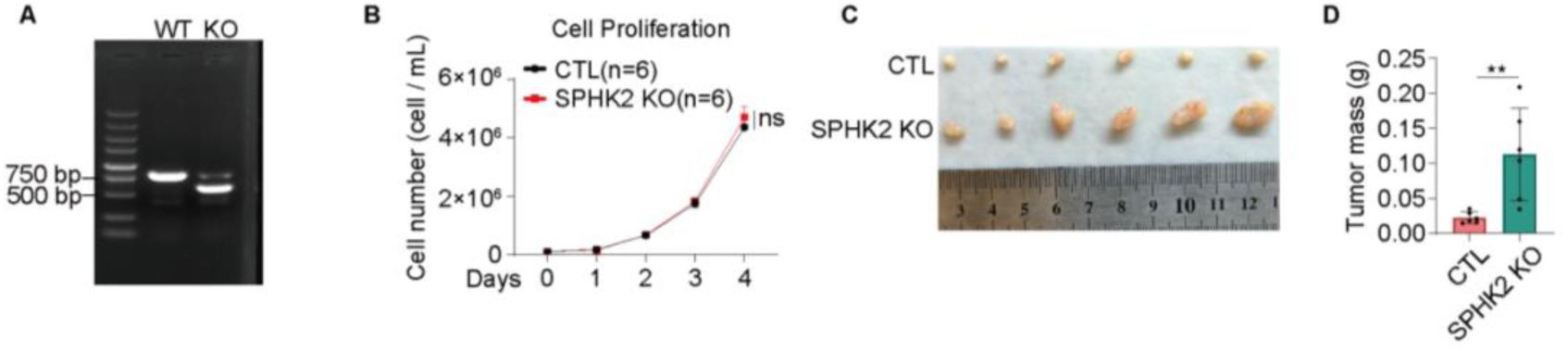
Tumor mitochondria do not affect the activation and exhaustion of CD8 cells **A**, Validation of SPHK2 knockout in MC38-Cox8a-GFP cells. **B**, Proliferation of SPHK2 knock out MC38 cell. **C-D**, Representative tumor image (**C**) and tumor weight (**D**) of SPHK2-KO MC38 subcutaneous tumors. None Significance: ns (*P* > 0.05), \**P* < 0.05, \*\**P* < 0.01, \*\*\**P* < 0.001, \*\*\*\**P* < 0.0001 (unpaired Student’s t-test).

